# Striatal low-threshold spiking interneurons locally gate dopamine during learning

**DOI:** 10.1101/2020.08.03.235044

**Authors:** Elizabeth N. Holly, M. Felicia Davatolhagh, Rodrigo A. España, Marc V. Fuccillo

**Affiliations:** Department of Neuroscience, University of Pennsylvania, Philadelphia, PA; Neuroscience Graduate Group, Perelman School of Medicine, University of Pennsylvania, Philadelphia, PA; Department of Neurobiology and Anatomy, Drexel University College of Medicine, Philadelphia, PA

## Abstract

Low-threshold spiking interneurons (LTSIs) in the dorsomedial striatum are potent modulators of goal-directed learning. Here, we uncover a novel function for LTSIs in locally and directly gating striatal dopamine, using in vitro fast scan cyclic voltammetry as well as in vivo GRAB-DA sensor imaging and pharmacology during operant learning. We demonstrate that LTSIs, acting via GABAB signaling, attenuate dopamine release, thereby serving as local coordinators of striatal plasticity.

---

The dorsomedial striatum (DMS) is a central hub supporting goal-directed learning and performance. However, it remains unclear how this structure locally integrates widespread cortical, thalamic, and limbic inputs to orchestrate behavior. Increasingly, evidence points to a key role of local inhibitory networks in coordinating the flow of striatal inputs with the output of striatal projection neurons to drive behavior^1^. Recently, we demonstrated that one subtype of GABAergic interneuron in the DMS, low-threshold spiking interneurons (LTSIs), plays an important function in initial goal-directed learning^2^. As a population, LTSIs exhibit robust reward-circumscribed activity that is downmodulated across operant learning. Inhibiting LTSIs during reward retrieval accelerates learning, while maintaining LTSI activity during this reward period slows learning. As LTSIs exert state-dependent control over DMS spiny projection neuron (SPN) activity and functional output^3^, dynamic LTSI downmodulation during reward learning may promote strengthening of corticostriatal circuits, permitting the transition from non-focused, highly variable motor output to stable, efficient motor behavior.

Dopamine signaling is a major reward-related neuromodulatory system driving goal-directed learning and performance. Striatal dopamine serves as a teaching signal^4^, facilitating synaptic plasticity^5^ and invigorating motor behavior^6^. This functional diversity is supported by an array of regulatory mechanisms to control dopamine signals across multiple temporal and spatial regimes. Importantly, local microcircuitry gates dopamine at the terminals, regulating the magnitude, spread, timing, and duration of dopamine signals^7^. A range of local homosynaptic and heterosynaptic mechanisms directly regulate dopamine at striatal terminals^7^, including striatal GABA tone^8–10^. Given the dynamic activity of LTSIs during reward retrieval, we hypothesized that LTSIs might provide a novel GABAergic mechanism of local dopamine regulation in the striatum with significant implications for learning.

To probe for initial evidence of LTSI-dopamine interactions, we performed in vivo microdialysis of extracellular striatal dopamine. Inhibition of striatal LTSIs via overexpression of Kir2.1, an inwardly rectifying potassium channel, significantly augmented extracellular dopamine evoked by a low dose of amphetamine (Supplemental Figure 1). While this evidence supports an interaction between LTSIs and dopaminergic transmission, multiple potential mechanisms could generate these effects: (1) direct dopamine axon inhibition, (2) disinhibition of local cholinergic interneurons (ChINs), which strongly regulate dopamine^7,11^, and/or (3) disinhibition of local spiny projection neurons (SPNs), which can regulate dopamine via recurrent circuitry^12^.

We were intrigued by the possibility that LTSIs could directly innervate dopamine axons to locally regulate release. Anatomical evidence suggests direct axo-axonal interactions occur between LTSIs and dopamine processes - synaptophysin-labeled LTSI synapses co-localize with tyrosine hydroxylase immunoreactive fibers in close proximity to synaptophysin-labeled dopaminergic terminals (Supplemental Figure 2). We directly probed for functional interactions between LTSIs and dopamine axons using ex vivo fast scan cyclic voltammetry. We eliminated potential contributions of ChINs by optogenetically stimulating dopamine terminals and performing all experiments in the presence of nicotinic and muscarinic acetylcholine antagonists. We eliminated downstream effects of LTSI-SPN interactions by performing experiments in acute coronal striatal slices, which disrupts SPN projections to the midbrain. LTSIs are tonically active in slice, so we first tested the effects of LTSI inhibition on ChR2-mediated optically evoked dopamine (oDA; Figure 1a-c). Halorhodopsin-mediated LTSI inhibition augmented oDA when dopamine axons were stimulated in the middle of both 4s (mirroring our prior in vivo manipulation) and brief 400ms LTSI inhibition (Figure 1c, ‘Mid’). We tested the temporal precision of LTSI-DA control, noting a similar oDA augmentation when LTSI inhibition terminated 500ms prior to dopamine axon stimulation (Figure 1c, ‘Delayed’). This prolonged effect of LTSI inhibition suggests longer lasting metabotropic as opposed to ionotropic mechanisms of regulation.

**Figure 1.**
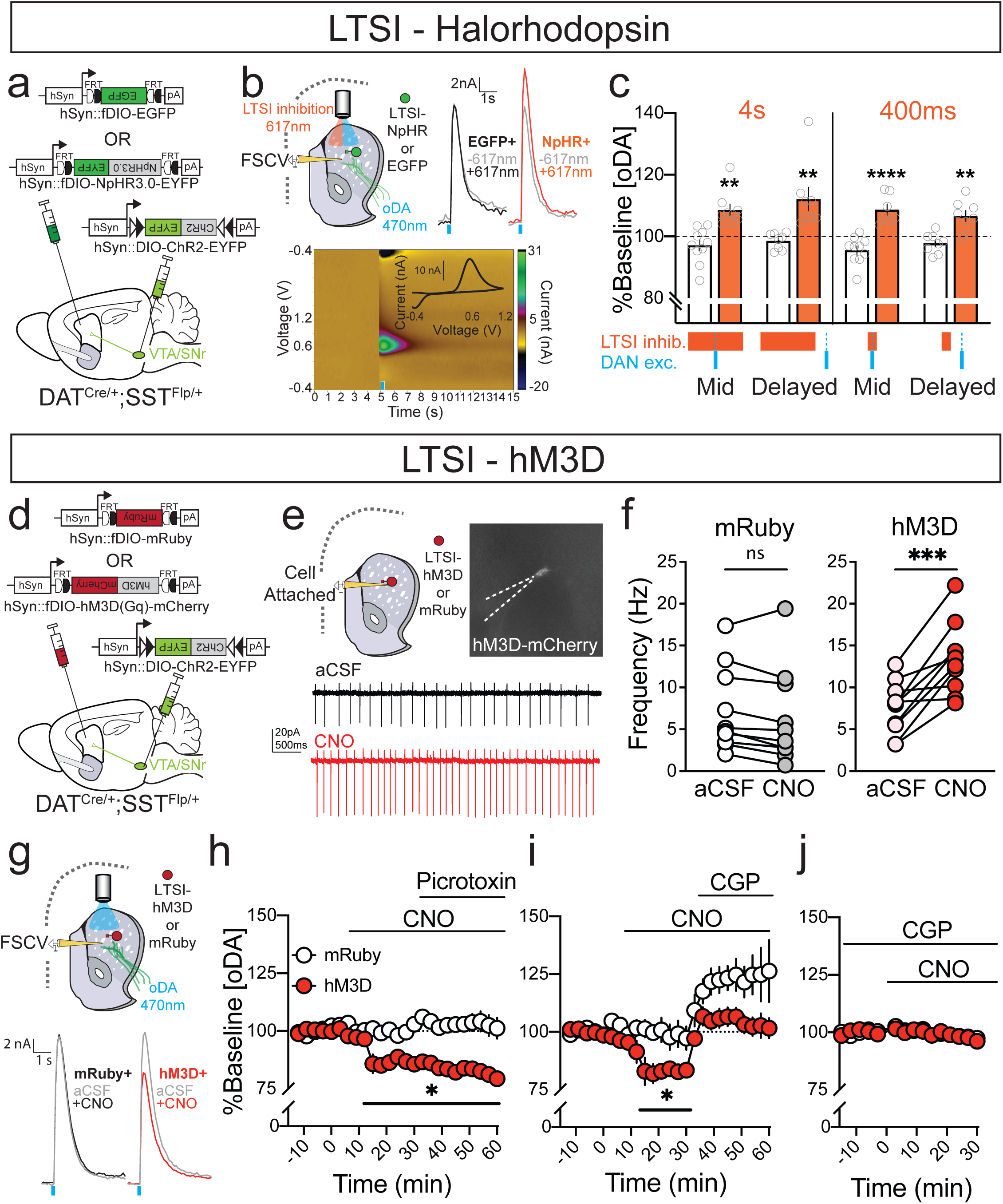
LTSIs attenuate optogenetically evoked dopamine via GABA_B_ signaling. (a) Experimental design for optogenetically driven LTSI inhibition and dopamine terminal excitation. (b) Recording schematic (top left), sample traces (top right) and voltammogram (bottom) for acute slice fast scan cyclic voltammetry (FSCV) experiments testing the effect of LTSI inhibition on optogenetically-evoked dopamine (oDA). (c) Percent change from baseline oDA in striatal slices with LTSIs expressing EGFP (white) or halorhodopsin (orange). oDA was evoked by 2ms 470nm light stimulation in the middle (‘Mid’) or 500ms after termination (‘Delayed’) of 4s (left) or 400ms (right) 617nm light stimulation. **p<0.01, **** p<0.00001 vs LTSI-EGFP control in same stimulation condition. Data expressed as mean ± SEM, with individual values shown in gray. (d) Experimental design for chemogenetic LTSI excitation and optogenetic dopamine terminal excitation. (e) Recording schematic and sample traces for cell-attached electrophysiological recordings. (f) Spontaneous firing frequency (Hz) in mRuby+ (left; n=10 cells / 4 mice) and hM3D+ (right; n=10 cells / 4 mice) LTSIs in the presence of aCSF and clozapine-N-oxide (CNO, 10μM). **p<0.01 pairwise comparison of firing frequency. (g) Recording schematic and sample traces for acute slice FSCV experiments testing the effect of LTSI excitation on oDA. (h) Effects of GABA_A_ antagonism on oDA suppression caused by chemogenetic LTSI excitation. CNO (10μM) was applied after a stable baseline (<10% variability in 5 consecutive oDA stimulations), and picrotoxin (100μM) was added 30min later on slices where LTSIs expressed mRuby (n=7) or hM3D (n=8). *p<0.05 vs baseline. Data expressed as mean ± SEM. (I,j) Effects of GABA_B_ antagonism on oDA suppression caused by chemogenetic LTSI excitation. The GABA_B_ antagonist CGP55845 (2μM) was applied (i) 30min after CNO or (j) was present in the recording solution during baseline sample collection prior to CNO application (right). *p<0.05 vs baseline. Data expressed as mean ± SEM. See Supplemental Table 1 for detailed statistics.

Growing evidence points to a clear role of tonic GABA signaling locally modulating dopamine release in the striatum via both GABA_A_ and GABA_B_ transmission^8–10^. As LTSIs are tonically active in striatal slices, they are a compelling candidate source for this modulatory GABAergic tone. To mechanistically probe how LTSIs gate oDA, we next employed chemogenetic activation of hM3D-Gq to increase LTSI tonic firing (Figure 1d-f). Supporting a bidirectional role of LTSI-dopamine interactions, increasing LTSI activity suppressed oDA. This suppression was not affected by GABA_A_ antagonism (Figure 1h). In contrast, GABA_B_ antagonism inhibited the LTSI excitation-induced oDA suppression (Figure 1i), and prevented oDA suppression when applied prior to LTSI excitation (Figure 1j). Together, these experiments suggest that LTSIs gate striatal dopamine release via GABA_B_ locally and directly, independently of ChIN- or striatal loop-mediated mechanisms.

We next interrogated whether LTSI inhibition also alters dopamine signaling in vivo during learning. To monitor striatal dopamine with sub-second resolution, we imaged the GRAB-DA dopamine sensor^13^ as mice acquired a self-initiated operant task^2^ (Figure 2a-c). We modeled learning as a sigmoidal function of accuracy, with the active learning period defined as trials where accuracy rapidly increased (see Methods; Figure 2d). Replicating and extending our prior work^2^, we show that Kir2.1-mediated LTSI inhibition in the DMS accelerates learning (Figure 2e, Supplemental Figure 3a), with increased accuracy during the period of active learning (Figure 2f) driven by fewer omissions (Supplemental Figure 3c) but not incorrect responses (Supplemental Figure 3d). Notably, LTSI inhibition did not alter the duration of the pre-learning period, but rather the rate at which learning occurred (Figure 2e). While LTSI inhibition increased response rate and decreased lever press latency, initiation and reward retrieval latencies were unaffected (Supplemental Figure 3e-h).

**Figure 2.**
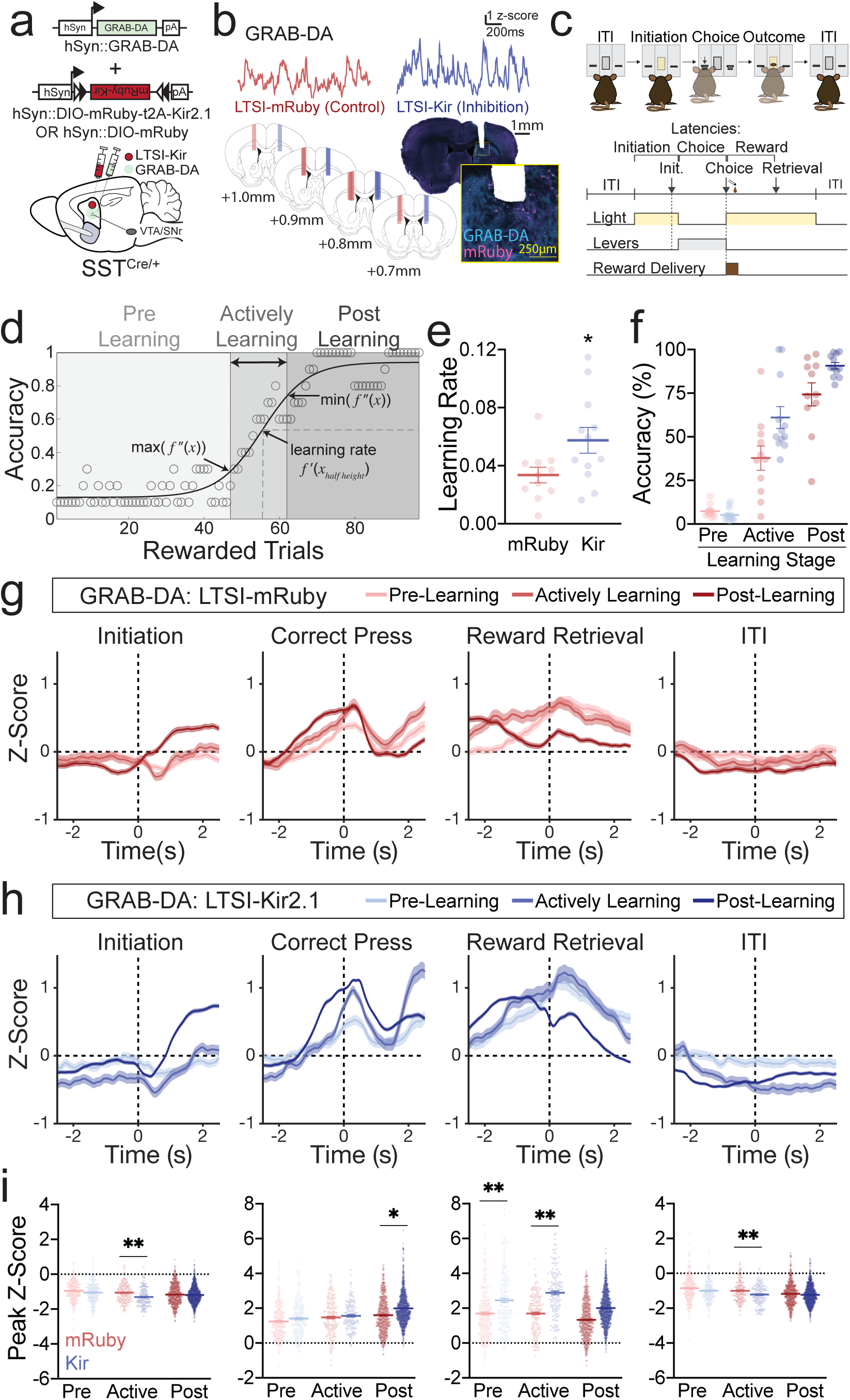
LTSI inhibition amplifies dopamine signaling and accelerates operant learning. (a) Experimental design for recording dopamine signals with or without LTSI inhibition during operant learning. (b) Sample GRAB-DA dopamine sensor traces (top) and fiberoptic cannula placement (bottom). (c) Self-initiated operant task. (d) Sigmoidal modeling of learning. For each trial, accuracy was calculated as the number of correct presses in the preceding 10 initiated trials. Only rewarded trials are depicted for simplification. The inflection points of the sigmoidal function (maximum and minimum values of the second derivative) were used to bin trials into ‘pre-learning’, ‘actively learning’, and ‘post-learning’ periods. The learning rate was defined as the instantaneous slope at the half height of the function. (e) Learning rate in mice with LTSI inhibition (Kir, n=12) or control (mRuby, n=11). *p<0.05 vs LTSI-mRuby control. (f) Accuracy in pre-learning, active learning, and post-learning periods. Two-way repeated measures ANOVA: virus p<0.05, learning stage p<0.001, interaction p<0.01. (g) Peri-event temporal histograms (PETHs) for initiations, correct presses, reward retrievals, and ITIs of pre-learning, actively learning, and post-learning trials for LTSI-mRuby control mice (n=11). Dashed vertical lines at time 0 indicate the timestamp of the behavioral event. (h) PETHs for LTSI-Kir mice (n=12). (i) Peak Z-scores (minima for initiation and ITI, maxima for correct press and reward retrieval) in the 1s window surrounding each behavioral event. *p<0.05, ***p<0.001 vs mRuby control at same learning stage. Lines in dot plots represent means; all PETH data represented as mean ± SEM. See Supplemental Table 1 for detailed statistics.

Consistent with our microdialysis and FSCV results, LTSI inhibition amplified the overall height and frequency of GRAB-DA dopamine peak events (Supplemental Figure 3b). We next explored whether LTSI inhibition selectively affected dopamine signals during discrete behavioral events or learning stages. The operant task structure allows for dissection of signals in response to trial initiation, lever press, reward retrieval, and an inter-trial interval (Figure 2c). By aligning dopamine signals to these specific behavioral events (Figure 2g-i, Supplemental Figure 3i), we revealed dynamic changes in dopamine signals across learning stages.

In pre-learning trials (those on the sigmoidal function prior to the rapid increase in accuracy), we observed dopamine signals circumscribed to the correct lever press and retrieval of the reward (Figure 2g). As learning proceeded, the dopamine signal connected to the correct lever press grew, while the dopamine signal aligned to the reward retrieval decreased (Figure 2g). LTSI inhibition did not alter the general progression of dopamine signals across learning, but rather augmented select signals at specific learning stages (Figure 2h,i). Prior to learning, LTSI inhibition selectively amplified dopamine signals during reward retrieval, while transients during other behavioral epochs remained unaffected. During and after task acquisition, LTSI inhibition amplified both choice- and reward-related dopamine signals (Figure 2h,i).

To gain further insight into how movement direction is integrated into these dopamine signals, we performed separate analyses of mice trained to press the lever contralateral (Supplemental Figure 4) or ipsilateral (Supplemental Figure 5) to their photometry implant. Consistent with prior reports^14,15^, we observed stronger striatal dopamine signals during contralateral movements compared to ipsilateral movements. Dopamine signals grew across learning as mice approached and pressed the lever contralateral to their photometry implant (Supplemental Figure 4a-c). In contrast, in mice trained to press the lever ipsilateral to the photometry implant dopamine signals did not peak until after the lever press, as mice were initiating a contralateral movement back towards the reward magazine (Supplemental Figure 5a-c). LTSI inhibition amplified dopamine signals in response to contralateral movement towards the magazine only in early stages of learning (Supplemental Figure 5d-e), while contralateral movements towards the lever were amplified in later stages of learning (Supplemental Figure 4d-e).

Overall, we find that as mice learn a goal-directed task, dopamine signals shift from reward-oriented to contralateral movement-oriented. Furthermore, LTSI inhibition amplifies dopamine signals in response to reward, particularly in early learning, which may contribute to accelerated task acquisition. To test this, we combined viral LTSI manipulation with local microinjection of aripiprazole, a dopamine D_2_ partial agonist. Aripiprazole increases dopamine synthesis when basal dopamine is low, while decreasing dopamine synthesis when basal dopamine is higher^16,17^. We hypothesized aripiprazole would stabilize striatal dopamine levels between LTSI-inhibited and LTSI-control mice, thereby suppressing an enhancement in learning rate mediated by LTSI inhibition. Microinjection of 100ng aripiprazole into the DMS prior to operant acquisition sessions (Figure 3a,b) blocked the effects of LTSI inhibition on operant learning rate (Figure 3c, Supplemental Figure 6). Importantly, these aripiprazole doses did not produce overt effects on overall motor activity as measured by operant task latencies (Supplemental Figure 6d-f) and rate of lever pressing (not shown) and as separately recorded in an open field (Figure 3d).

**Figure 3.**
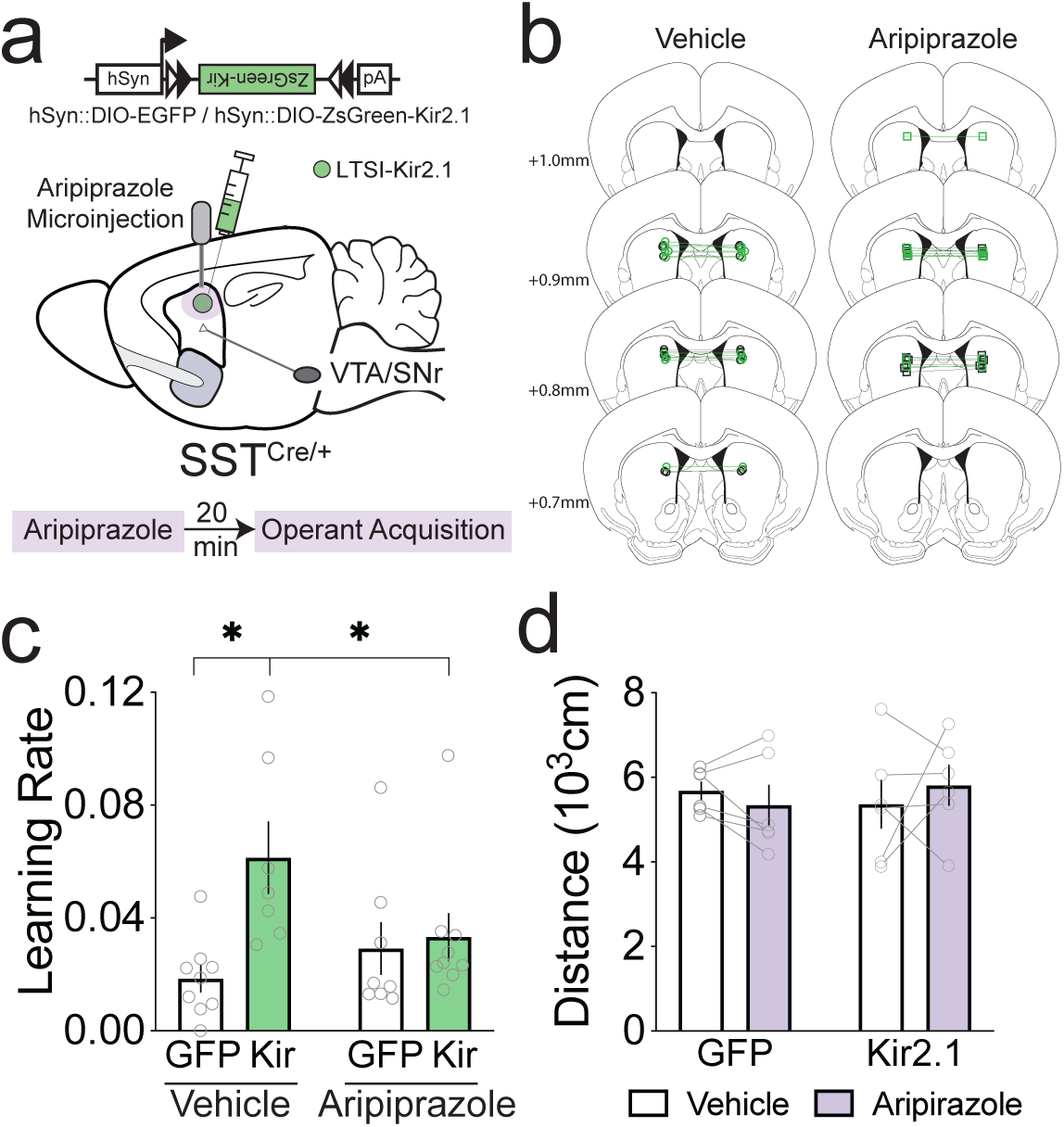
Intra-striatal dopamine D_2_ partial agonism prevents the effects of LTSI inhibition on learning. (a) Experimental design. Aripiprazole (100 ng/side) or vehicle was bilaterally microinjected into the dorsomedial striatum of mice with or without Kir-mediated LTSI inhibition prior to operant task acquisition sessions. (b) Placements of microinjector tips in the dorsomedial striatum. Vehicle (left): n=9 LTSI-EGFP (black circles), n=8 LTSI-Kir2.1 (green circles); Aripiprazole (right): n=8 LTSI-EGFP (black squares), n=9 LTSI-Kir2.1 (green squares). (c) Learning rate. P<0.05 vs Vehicle-Kir. (d) Distance traveled in an open field in a subset of LTSI-GFP (n=6) and LTSI-Kir (n=6) expressing mice one week after operant acquisition. All data represented as mean ± SEM with individual data shown. See Supplemental Table 1 for detailed statistics.

Taken together, these data demonstrate that LTSIs provide a novel mechanism for local modulation of dopaminergic signalling, acting via GABA_B_ signaling to gate synaptically released dopamine. This dynamic regulation occurs both in slice and *in vivo*, and underlies the effects of LTSI manipulations on operant learning. Due to the biophysical constraints of lengthy and highly-branched dopamine axons, local regulatory mechanisms are necessary to further refine striatal dopamine output^18^. By virtue of their unique reward-associated moudlation^2^, LTSIs are ideally situated to control the timing and magnitude of dopamine signals during learning. In addition to directly gating striatal dopamine, LTSIs further participate in this local microenvironment through strong inhibitory control over ChINs^19^, which also directly and strongly modulate striatal dopamine^11^. As both LTSIs and ChINs are dynamically modulated by discrete components of reward learning and performance, future work should delineate how these interneurons work in concert to shape dopamine signals during ongoing goal-directed behavior.

Corticostriatal connectivity drives action selection and performance, and plasticity in these circuits is critical for motor control and learning. We suggest that LTSIs act as a local coordinator of corticostriatal plasticity, owing to their combined local modulation of dopamine and dendritic inhibitory functions^19^. During early learning, dopamine solidifies corticostriatal eligibility traces, facilitating long term plasticity^20^ and strengthening the association between action selection and outcome. As we demonstrate, striatal LTSI activity can gate this dopaminergic facilitation, accelerating operant acquisition. As learning progresses, LTSI amplification of dopamine signaling may invigorate movement through direct involvement in action selection or execution^14^. In parallel to local dopamine modulation, LTSIs control the flow of cortical input to the striatum via state-dependent feedforward inhibitory actions on SPN distal dendrites^3,19^. Removing the LTSI brake on both striatal dopamine and distal dendritic compartments in a coordinated manner likely facilitates synaptic plasticity, thereby strengthening action-outcome associations and promoting future goal-directed behavior.

## Acknowledgements

This work was supported by NIMH F32 MH114506 to ENH, Howard Hughes Medical Institute Gilliam Fellowship to MFD, NIMH R00MH099243, R01 MH118369, and Whitehall Foundation grants to MVF. We thank William Doyon and John Dani for HPLC use for microdialysis experiments, Deborah Kwon and Zhaolan (Joe) Zhou for assistance with the open field experiment, and Rodrigo España, Emily Black, Jessica Shaw, Jung Hoon Shin, and Veronica Alvarez for assistance establishing fast scan cyclic voltammetry methodology. We also thank Nathan Henderson for initial confocal imaging pilot work, and Andrea Stout and the Cell and Developmental Biology Microscopy core at the Perelman School of Medicine for the confocal imaging presented in this paper. Finally, we thank Patrick Rothwell for constructive feedback on the manuscript draft.

## Author Contributions

Conceptualization, ENH and MVF; Methodology ENH, RAE, MVF; Formal Analysis ENH; Investigation ENH, MFD; Writing – Original Draft, ENH; Writing – Review and Editing, ENH, MFD, RAE, MVF; Funding Acquisition, ENH, MVF.

## Methods

### Contact for reagent and resource sharing

Further information and requests for resources should be directed to and will be fulfilled by Marc Fuccillo. All relevant code can be found the Fuccillo-lab github site.

### Experimental model and subject details

All mice (SST-IRES-Cre, Jackson stock number 013044, RRID:IMSR_JAX:013044; SST-IRES-Flp, Jackson stock number 028579, RRID:IMSR_ JAX:028579; DAT-IRES-Cre, Jackson stock number 006660, RRID:ISMR_JAX:006660) were bred in house. Prior to experimental manipulation, mice were group housed with littermates on a 12:12 light-dark cycle and provided ad libitum food and water. Unless otherwise noted, all experiments were conducted on naïve adult male mice, which were randomly assigned to experimental groups. After surgical implantation of optical cannulas, mice in dopamine sensor photometry experiments were individually housed. All experiments were conducted in accordance with the National Institutes of Health Guidelines for the Use of Animals, and all procedures approved by the Institutional Animal Care and Use Committee of the University of Pennsylvania (protocol 805643). Sample sizes are detailed in figure legends and Supplemental Table 1.

### General stereotaxic surgery and viral injection methodology

Specific details regarding animals used, virus delivered, and implantation are provided below for individual experiments. As described previously^2^, viral injections and cannula implantations were performed on a stereotaxic frame (Kopf Instruments, Model 1900) under isoflurane anesthesia (1.5-2% + oxygen at 1 L/min) and body temperature maintained at 30°C throughout surgery (Harvard Apparatus, #50722F). Briefly, fur over the skull was removed with depilatory cream, and the skin cleaned with 70% isopropyl alcohol and betadine, after which a small anterior/posterior incision was made to expose the skull. Small (0.5 mm) holes were drilled above the target coordinates and a pulled glass needle was lowered into the injection site. DMS coordinates were AP: 0.85mm, ML: 1.25mm, DV: -2.85mm, unilaterally (dopamine sensor, anatomy) or bilaterally (all other experiments). VTA/SN coordinates were AP: -3.00mm, ML: ±0.65mm, DV-4.40mm. 500nl (DMS) or 1000nl (VTA/SN) of specific adeno-associated virus (see individual experimental details below) was infused at 125 nl/min using a microinfusion pump (Harvard Apparatus, #70-3007), and the injection needle was removed 10 min after termination of viral infusion. For implantation surgeries (microdialysis, dopamine sensor, microinjection), 2 small screws were secured into the skull. The implant was lowered into the injection site and held with dental cement (Den-Mat, Geristore A and B). Mice were given a minimum of 7d (microdialysis, microinjection) or 3 weeks (all other experiments) to recover from surgery prior to subsequent experimental testing.

### Histological verification of viral expression and implant placement

After the completion of behavioral experiments, mice were deeply anesthetized with i.p. pentobarbital (Nembutal, 50 mg/ml solution) and transcardially perfused with 4% formalin/PBS followed by PBS. Following overnight storage in 4% formalin, brains were stored in 30% sucrose until fully saturated, then sectioned via vibratome (Vibratome, Model 1000plus) or cryostat (Leica Microsystems, Model CM3050S). Viral expression and implant placements were imaged on a standard epifluorescent microscope (Olympus, BX63) under 4X (Olympus, 0.16NA) or 10x (Olympus, 0.4NA) objectives.

### *In vivo* microdialysis

The effects of LTSI inhibition on extracellular striatal dopamine were investigated using in vivo microdialysis. SSTCre/+ mice (n=4) were injected with AAVDJ.EF1α.DIO.ZsGreen-T2A-Kir2.1 in one striatal hemisphere and AAV1.CAG.DIO.GFP in the other striatal hemisphere. Microdialysis guide cannulae (3mm length; Synaptech, S-3000) were aimed at the injection site and implanted at a 15° angle (AP +0.85, ML ±1.99, DV-2.31).

Mice underwent *in vivo* microdialysis after 7-10d recovery from surgery. On the night before sample collection, a microdialysis probe with 1mm active polyacrylonitrile membrane (Synaptech, S-3010) was lowered into each guide. The probe was perfused with artificial CSF (aCSF, 149mM NaCl, 2.8mM KCl, 1.6mM CaCl2, 1.2mM MgCl2, 0.2mM ascorbic acid, 5.4mM D-Glucose) at a flow rate of 0.5μl/min. The following morning, the flow rate was increased to 2.0μl/min 2h prior to sample collection. Samples were collected by hand into 0.2 ml PCR tubes every 10min and stored on dry ice until the end of sample collection. After 5 baseline samples, mice were injected with saline (i.p.), followed 20 min later by d-amphetamine (1.0 mg/kg, i.p.). Samples were collected for 90 min following d-amphetamine administration. Following sample collection, mice perfused and probe placements and viral expression were verified as described above.

Dialysate was stored at -80°C until dopamine was analyzed by high performance liquid chromatography (HPLC) as described previously^21^. Briefly, mobile phase (4.0mM citric acid, 3.3mM sodium dodecyl sulfate, 100mM NaH2PO4, 0.3mM ethylenediaminetetraacetic acid, 15% acetonitrile, 5% methanol) was pumped (ThermoScientific, Model 582) through an HR-3.2 x 80mm column (3μm particle size, ThermoScientific) connected to a Coulochem II detector. An autosampler (ThermoScientific, Model 542) mixed 9.5μl dialysate with ascorbic oxidase (Sigma-Aldrich. EC 1.10.3.3; 162 units/mg) prior to injection, and dopamine signals acquired with 501 chromatography and Chromeleon Software (ThermoScientific). Dopamine concentration was quantified by comparing peak area to external standards (0-2.5 nM).

### Immunohistochemistry and confocal microscopy

To determine whether LTSIs make synaptic connections in close proximity to dopamine fibers, we unilaterally injected SST-Flp/+;DAT-Cre/+ mice (n=3) with AAVDJ.EF1α.fDIO.Synaptophysin-mRuby2 in the DMS to label LTSI terminals, and AAVDJ.EF1α.DIO. Synaptophysin-EGFP in the VTA/SN to label dopamine terminals. After 3-6 weeks of viral expression, mice were transcardially perfused with 4% formalin followed by PBS, and 50μm slices sectioned on a vibratome (Vibratome, Model 1000plus). Immunohistochemistry for tyrosine hydroxylase (TH) was performed to visualize the full extent of dopaminergic processes. Free floating sections were permeabilized in 0.6% Triton X-100 and blocked in 3% normal goat serum (NGS) in PBS for 1h. Primary antibody (mouse anti-TH, 1:4000, Immunostar #22941) was incubated overnight in 1% NGS and 0.2% Triton X-100 in PBS, followed by 2h incubation in secondary antibody (goat anti-mouse Alexa 647, 1:200, Invitrogen A-21236). Slices were then mounted and scanned on a Leica TCS SP8 STED 3X confocal with white light laser and imaged with a 40X/1,3 NA oil immersion objective. Synpatophysin-GFP puncta were excited by 490nm laser wavelength and detected with 500-545nm emission detection filter, Synpatophysin-mRuby puncta were excited by 555nm laser wavelength with 571-629 emission detection filter, and Alexa647 labeled TH fibers excited by 640nm laser wavelength with 652-746nm emission detection filter. To reduce contributions from autofluorescence and increase specificity of detection, time-gated detection using a HyD detector and time window of 0.4-6.5ns was used in all channels.

### Fast scan cyclic voltammetry (FSCV)

Optically evoked dopamine ([oDA]) was measured using fast scan cyclic voltammetry in acute striatal slices. For all FSCV experiments, SST-Flp/+;DAT-Cre/+ mice were injected with AAV5.hSyn.DIO.hChR2(H134R)-EFYP into the VTA/SN for optogenetic control of dopamine terminals. For LTSI optogenetic inhibition experiments (n=12, Fig1a-c), one striatal hemisphere was injected with AAVDJ.EF1α.fDIO.eNpHR3.0-EYFP and the other with AAVDJ.EF1α.fDIO-EYFP. For LTSI chemogenetic excitation experiments (n=16, Fig1d-i), one striatal hemisphere was injected with AAVDJ.hSyn. fDIO.hM3DGq-mCherry and the other with AAVDJ. EF1α.fDIO.mRuby2. Virus was allowed to express for at least 4 weeks prior to FSCV recordings.

To obtain acute striatal slices, mice were anesthetized with isoflurane and transcardially perfused with ice-cold sucrose cutting solution (225mM sucrose, 13.9mM NaCl, 26.2mM NaHCO_3_, 1mM NaH_2_PO_4_, 1.25mM glucose, 2.5mM KCl, 0.1mM CaCl_2_, 4.9mM MgCl_2_). The brain was removed, hemispheres bisected, and coronally sectioned (300μm) on a vibratome (Leica, Model VT1200s). Slices were incubated at 32°C for 15min in oxygenated (95% O_2_, 5% CO_2_) aCSF (124mM NaCl, 26.2mM NaHCO_3_, 1mM NaH_2_PO_4_, 10mM glucose, 2.5mM KCl, 2.5mM CaCl_2_, 1.3mM MgCl_2_, 0.4mM ascorbic acid), followed by at least 1h incubation at room temperature (20-22°C) prior to recordings.

For recording, slices were placed in a recording chamber, fully submerged in oxygenated aCSF at a flow rate of 1.4-1.6 ml/min, maintained at 30-32°C. All experiments were conducted in the presence of 1μM DHβE and 1μM scopolamine to preclude any possible effects of LTSI manipulation on cholinergic transmission. Carbon fiber electrodes (Kation Scientific, #E1011-20mod CarboStar1, custom 200μm tip length, 7μm diameter) were conditioned at 60Hz for 20 min in aCSF prior to first use. Carbon fiber electrodes were lowered 60-80μm into the DMS at a 20° angle.

To optogenetically evoke dopamine release (oDA), a 2ms pulse of 470nm light was illuminated through the 40x objective (Olympus, 0.8NA water immersion). Pulses were delivered every 3min, which allowed for stable release over several hours. A light intensity that elicited approximately 50% maximal [oDA] was determined for each recording location and used for experimental stimulation. A stable baseline (<10% variability in [oDA] over 5 consecutive samples) was established prior to LTSI manipulations. The scanning voltage was a triangular waveform (−0.4 to +1.2V vs Ag/AgCl), with a scan rate of 400 V/s every 100ms using a voltammeter (Dagan Corp., CHEM-CLAMP). Raw traces of oDA were analyzed using the Demon Voltammetry software package^22^.

#### Halorhodopsin-mediated LTSI inhibition

After a stable baseline was established (<10% variability in [oDA] across 5 consecutive samples), the effects of optogenetic LTSI inhibition were probed. Slices were illuminated with 617nm light (0.9 mW/mm^2^) through the 40X objective in one of four conditions: (1) 4s 617nm illumination, with oDA stimulation at 2s, (2) 4s 617nm illumination, with oDA stimulation 500ms after 617nm termination, (3) 400ms 617nm illumination, with oDA stimulation at 200ms, (4) 400ms 617nm illumination, with oDA stimulation 500ms after 617nm termination. Five [oDA] measurements were collected and averaged, and data expressed as percent change from the average of the five baseline [oDA] measurements.

#### Chemogenetic-mediated LTSI excitation

The effects of LTSI excitation on oDA were probed using chemogenetic activation of hM3D-Gq expressed in LTSIs. After a stable baseline was established, clozapine-n-oxide (CNO, 10μM) was added to the recording solution. The effects of GABA_A_ and GABA_B_ signaling were tested in three separate experiments. After 30min of CNO application, the GABA_A_ antagonist picrotoxin (100μM) or GABA_B_ antagonist CGP 55845 (2μM) was applied and [oDA] recorded for another 30min. In a separate experiment, CGP 55845 (2μM) was present during baseline sample collection, prior to application of CNO. Data were expressed as percent change from the mean of the last 5 baseline samples.

### Acute slice electrophysiology

Cell-attached recordings were used to validate chemogenetic-mediated increases in LTSI activity in slice. One striatal hemisphere of SST-Flp/+;DAT-Cre/+ mice (n=4) was injected with AAVDJ.hSyn.fDIO. mRuby2 and the other striatal hemisphere injected with AAVDJ.hSyn.fDIO.hM3DGq.mCherry.

Our general electrophysiology procedures have been described previously^2^. Briefly, mice were anesthetized with isoflurane and transcardially perfused with ice-cold aCSF (124 mM NaCl, 1.2 mM NaH_2_PO_4_, 2.5 mM NaHCO_3_, 5 mM HEPES, 13 mM glucose, 1.3 mM MgSO_4_, 2.5 mM CaCl_2_). The brain was then quickly removed and coronally sectioned (250 μm) on a vibratome (Leica, Model VT1200s). Slices were then incubated at 32°C for 12-15min in an NMDG-based recovery solution (92 mM NMDG, 2.5 mM KCl, 1.2 mM NaH_2_PO_4_, 30 mM NaHCO_3_, 20 mM HEPES, 25 mM glucose, 5 mM sodium ascorbate, 2 mM thiourea, 3 mM sodium pyruvate, 10 mM MgSO_4_, 0.5 mM CaCl_2_), then transferred to room temperature (20-22°C) aCSF for at least 1h before recording. For recording, slices were placed in a recording chamber, fully submerged in oxygenated (95% O_2_, 5% CO_2_) aCSF at a flow rate of 1.4-1.6 mL/min, and maintained at 29-30°C.

Cell-attached recordings of mRuby+ (n=10) or hM3D-mCherry+ (n=10) cells were made with electrodes filled with aCSF. We determined LTSI firing frequency (Hz) in 5 min recordings under baseline aCSF conditions to 5 min recordings in the presence of CNO (10μM). Neuronal spiking was detected by Neuromatic (v 3.0, Jason Rothman), and firing frequency was calculated as the overall spiking over the recording time window. Recordings were sampled at 20kHz and filtered at 2.8kHz. Data acquisition was in Igor 6.32 (Wavemetrics) using Recording Artist (Rick Gerkin) and analyzed offline in Igor 7.

### Operant Task

Methods for our operant learning task have been described in detail previously^2^. Briefly, mice were food deprived to 85-90% of free feeding weight prior to behavioral training. Experiments were conducted in a modular operant chamber (Med Associates Inc, Model ENV307W, 21.59 x 18.08 x 12.7cm) equipped with a modified liquid reward magazine flanked by retractable levers on either side. Chocolate liquid reward (Nestlé Boost, 10μl) served as the positive reinforcer, delivered into the reward magazine by a pump (Med Associates Inc, Model PHM-100). Mice were first familiarized with the operant chambers in magazine training sessions, where 10μl reward was delivered at the onset of 10s magazine light illumination once per minute for 40min. These sessions continued for a minimum of 2 days until mice had fewer than 10 omissions (trials in which mice did not retrieve the reward within 10s of magazine light illumination).

Mice were then trained on a fixed ratio 1 (FR1) self-initiated two-choice operant task (Figure 2c). The task structured into four discrete phases: (1) Inter-trial interval (ITI) - all lights were off for 5s between each trial), (2) Initiation - magazine light illuminated, and nosepoke initiated the trial), (3) Choice - extension of both retractable levers for either 10s or until a lever was pressed. In the event of an omission (10s without press), levers were retracted and the trial ended, (4) Outcome - mice were randomly assigned a correct lever (left or right); pressing the correct lever resulted in 5s magazine light illumination and 10μl chocolate reward, ending the trial, while pressing the incorrect lever ended the trial.

#### Sigmoidal modeling of learning curves

Learning curves for each mouse were modeled by fitting trial accuracy over time to a sigmoidal function. For each trial, accuracy was defined as the percentage of rewarded trials in the previous ten initiated trials (including correct choice, incorrect choice, and omission). Accuracy over trials was then fit with sigm_ fit from the MATLAB Central File Exchange to the sigmoidal function

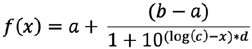

where *a* is the minimum, *b* is the maximum, *c* is the x value at half height of the function, and *d* is the slope. To next bin the trials into pre-learning, active learning, and post-learning periods, the maximum and minimum values of the second derivative of the function were used. These values are the inflection points of the curve, demarcating when the slope is changing direction. Learning rate was defined as the instantaneous slope of the trial at the half-height of the sigmoidal function (Figure 2d).

### Fiber photometry

Mice were unilaterally injected with AAV9.hSyn. GRAB-DA(v4.4) combined with either AAVDJ.EF1α. DIO.mRuby2 (n=11) or AAVDJ.EF1α.DIO.mRuby2-Kir2.1 (n=12) into the DMS and implanted with a 400μm diameter fiberoptic cannula (0.48NA, 3-5mm length, constructed in house). Viruses were allowed to express for at least 3 weeks until stable dopamine sensor transients were observed. Mice were then food deprived and underwent reward magazine training and operant training as described above while photometric dopamine signals were recorded.

#### Signal Collection

Fiber photometry was performed as described previously^2^. Mice were attached via an optical fiber (400μm core, 0.48NA; Doric Lenses), which was connected to a Doric 4-port minicube (FMC4, Doric Lenses). Dual color LED light (470nm for GRAB-DA stimulation, ThorLabs #MF470F3; 405nm for artifact control fluorescence, ThorLabs #MF405FP1) was delivered through the fiberoptic cannula into the brain at 10-30μW (ThorLabs, LED Driver Model DC4104). Photon emissions were passed through a dichroic mirror and 5000-550nm cult filter, then detected by a femtowatt silicon photoreceiver (Newport, Model 2141). Analog signals were demodulated and recorded with an RZ5 processor and Synapse Software (Tucker Davis Technologies). Prior to each recording session, 470nm light was passed through the patch cord for at least 4h to reduce autofluorescence.

Mice were connected to the patch cord with the LEDs on in the operant chambers for at least 10 min prior to experimental sessions to allow for habituation. All recording sessions began with a 10 min baseline period. Operant training sessions as described above were typically 60min, but allowed to extend longer (no more than 120min) if the mouse had started to acquire the task but was not yet exhibiting consistent accurate performance.

#### Signal Analysis

The demodulated 470nm signal was processed and analyzed with custom scripts written in Matlab (MathWorks, Version 2017b). Data analysis was adapted from our previously described methodology^2^, with some modifications made to optimize for dopamine signal analysis. Data were down-sampled to 40Hz then digitally filtered (filtfilt in Matlab). Over extended recording sessions, there may be steady decreases in baseline autofluorescence. In order to account for this, the 470nm signal at the end of the recording was baselined to zero, the data fit to a double exponential curve, and ΔF/F calculated as (F-F_0_)/F_0_. The z-score was calculated as the difference between the ΔF/F and mean ΔF/F for the recording session, divided by the standard deviation of the ΔF/F across the recording session.

Dopamine sensor peak events were calculated using custom peak detection scripts^2,23^. A 10s moving window was used for thresholding, where high amplitude events (local maxima greater than two median average deviations above the median of the moving window) were removed to calculate a new baseline moving median. Peaks were defined as events with local maxima greater than 3 median average deviations above this new baseline moving median. Peak amplitude was calculated as the difference between the peak maxima and the local median.

We also assessed dopamine sensor activity tied to discrete behavioral events using peri-event temporal histogram (PETH) analysis. The z-scored ΔF/F signal was aligned to time 0 for each behavioral timestamp (delivered by TTL signal from MedPC to Synapse software) and a the 2.5s before and after the timestamp were extracted. The peak z-score (minima for ITI and initiation, maxima for choice and reward retrieval signals) and location were extracted from the 1s window around the behavioral event and areas under the curve (AUC) calculated for each trace.

### Dopamine D_2_ partial agonist (aripiprazole) microinjection

SSTCre/+ mice were bilaterally injected with either AAVDJ.EF1α.DIO.ZsGreen-Kir2.1 (n=17) or AAV1. CAG.DIO-EGFP (n=17), and implanted with bilateral microinjection cannulae (Plastics1, C235G-3.0/SPC 4mm length). Dummy cannulae with 1mm protrusion (Plastics1, 235DC/SPC) were inserted in microinjector cannulae and held in place with a dust cap (Plastics1, 303DC/1). Mice were given 7d to recover before food deprivation and operant training.

#### Microinjection procedure

Microinjections occurred 20 min prior to magazine and operant training sessions. Dummy cannulae were replaced with microinfusion cannulae (Plastics1, C235I/SPC 5mm, 1mm projection beyond guide) connected by PE50 tubing to an infusion pump (Harvard Apparatus, 704506 Pump 11 Pico Plus Elite). Aripiprazole (100ng/side) or vehicle was administered in a volume of 250nl/side across 2min, and injectors left in place for an additional 1min to allow for diffusion from the injection site and prevent backflow. The dose for aripiprazole was selected based on prior research^24^. To acclimate mice to microinjection procedures, all mice received vehicle (see ‘Drugs’ below) infusions prior to the three magazine training sessions. Subsequently, mice received either aripiprazole (n=8 LTSI-GFP, n=9 LTSI-Kir) or vehicle (n=9 LTSI-GFP, n=8 LTSI-Kir) prior to each 1h operant training session. Mice continued operant training until they obtained 50 rewards within one session.

### Open Field

After completing operant training, a subset of mice (n=6 LTSI-GFP, n=6 LTSI-Kir) were tested in an open field to evaluate the effects of aripiprazole on general locomotor behavior. Mice were first habituated to the open field arena (15” x 15” box) placed directly underneath a ceiling-mounted camera in a one hour session. The periphery and dimensions of the arenas were defined in video tracking software (SmartScan 3.0) and total distance traveled was recorded and used for analysis. The next day, mice were randomly assigned to receive vehicle or aripiprazole microinfusion, after which they were placed in the open field arena. Recording began 20 min after the microinfusion, and distance traveled measured for 30 min. The following day, the procedure was repeated with mice receiving the opposite microinfusion.

### Drugs

Dihydro-β-erythroidine (DHβE) and aripiprazole were obtained from Tocris Bioscience; all other chemicals were obtained from Sigma-Aldrich. Aripipirazole was dissolved in 1μl glacial acetic acid, then brought to volume with deionized water and pH adjusted to 5.2 with NaOH. The vehicle was prepared in the same way, without the addition of aripiprazole. Aripiprazole and vehicle stocks were made the same day, aliquoted, and stored at -20°C. Stock solutions for drugs used in fast scan cyclic voltammetry were prepared in deionized water (100mM DHβE, 100mM scopolamine, 100mM CGP55845) or DMSO (100mM picrotoxin, 25mM clozapine-N-oxide), aliquoted and stored at -20°C.

### Statistical Analysis

General linear mixed models were performed with SAS (SAS Institute, University Edition), and all other statistical analyses were performed with Prism software (GraphPad, version 8). Detailed statistics and sample sizes for each figure panel can be found in Supplemental Table 1. Appropriate t-tests (paired and unpaired), ANOVAs (one-way, two-way, two-way repeated measures, three-way repeated measures), and general linear mixed models (GLMM) were perform as indicated in the results and supplemental table. ANOVAs/GLMMs with significant main effects/interactions were followed up with a priori driven post-hoc tests with Bonferroni corrections for multiple comparisons. Data that violated assumptions of normality were transformed as indicated in the supplemental table, and Geisser-Greenhouse corrections applied to data that violated assumptions of sphericity. Kenward-Roger corrections (KENWARDROGER2) were applied in GLMMs to correct degrees of freedom for fixed effects.

