## Supplemental Figures for "Striatal low-threshold spiking interneurons locally gate dopamine during learning"

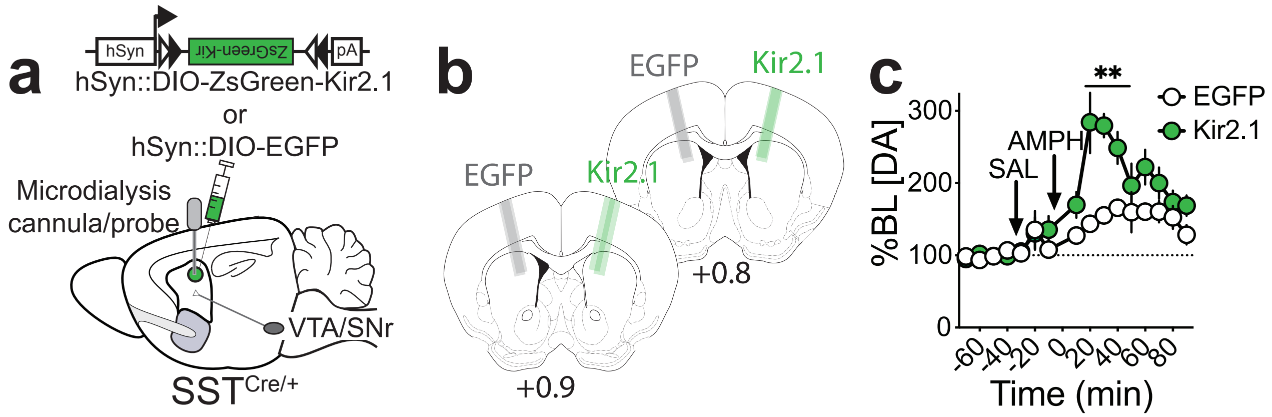


**Supplemental Figure 1. LTSI inhibition augments striatal dopamine in response to d-amphetamine challenge.** (a) Experimental design. (b) Probe placements for 4 mice expressing EGFP in the LTSIs of one hemisphere (gray) and Kir2.1 in LTSIs of the other hemisphere (green). (c) Percent change from baseline extracellular dopamine after saline (SAL, i.p.) and d-amphetamine (AMPH, 1.0 mg/kg, i.p). **p<0.01 vs baseline. Data represented as mean ± SEM. See Supplemental Table 1 for detailed statistics.

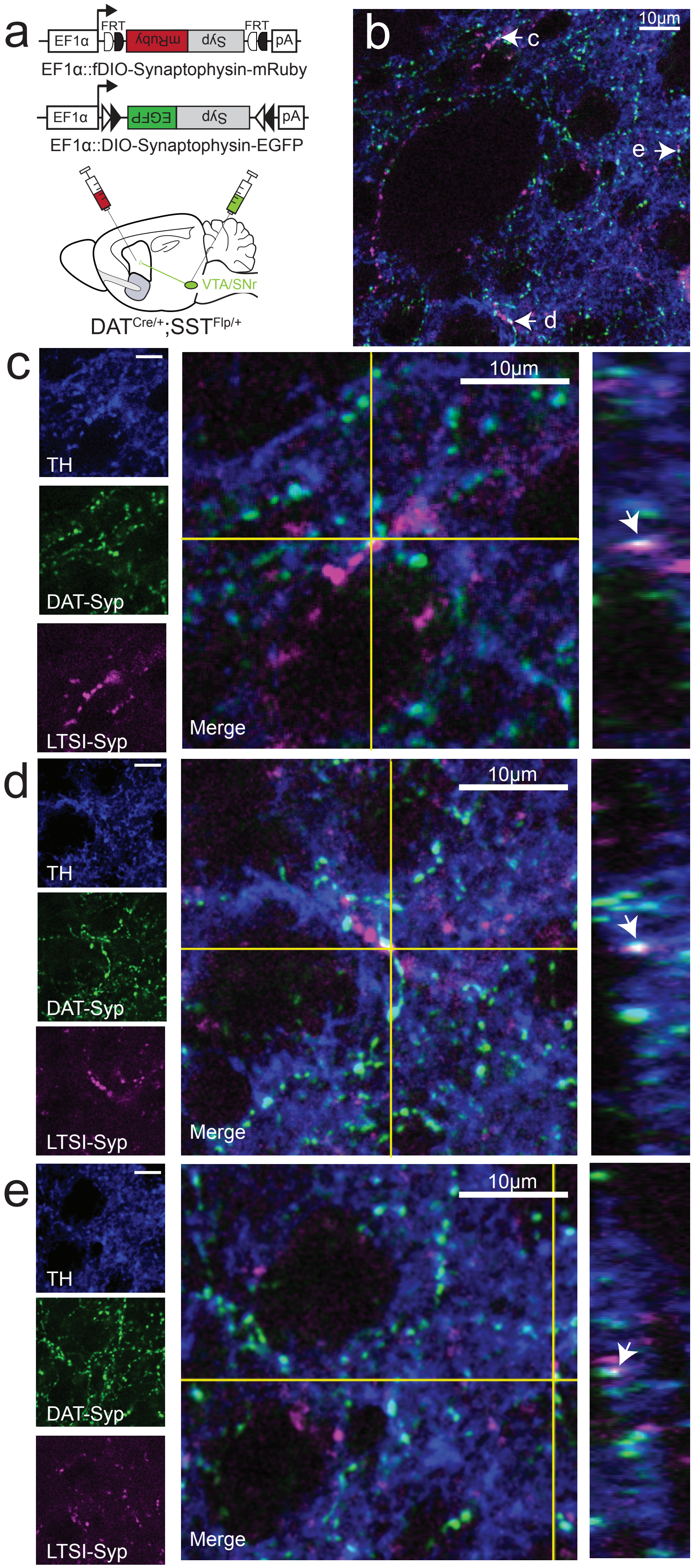


**Supplemental Figure 2. LTSIs synapse in close proximity to dopaminergic synapses.** (a) Experimental design. (b) Representative 40X images. White arrows indicate colocalizations between LTSI-Synaptophysin-mRuby (magenta), DAT-Synaptophysin-GFP (green), and tyrosine hydroxylase (TH) immunoreactive fibers (blue). (c,d) Enlarged regions with co-localizations labeled in (b). Orthogonal YZ view shown on right, with arrow pointing to co-localization.

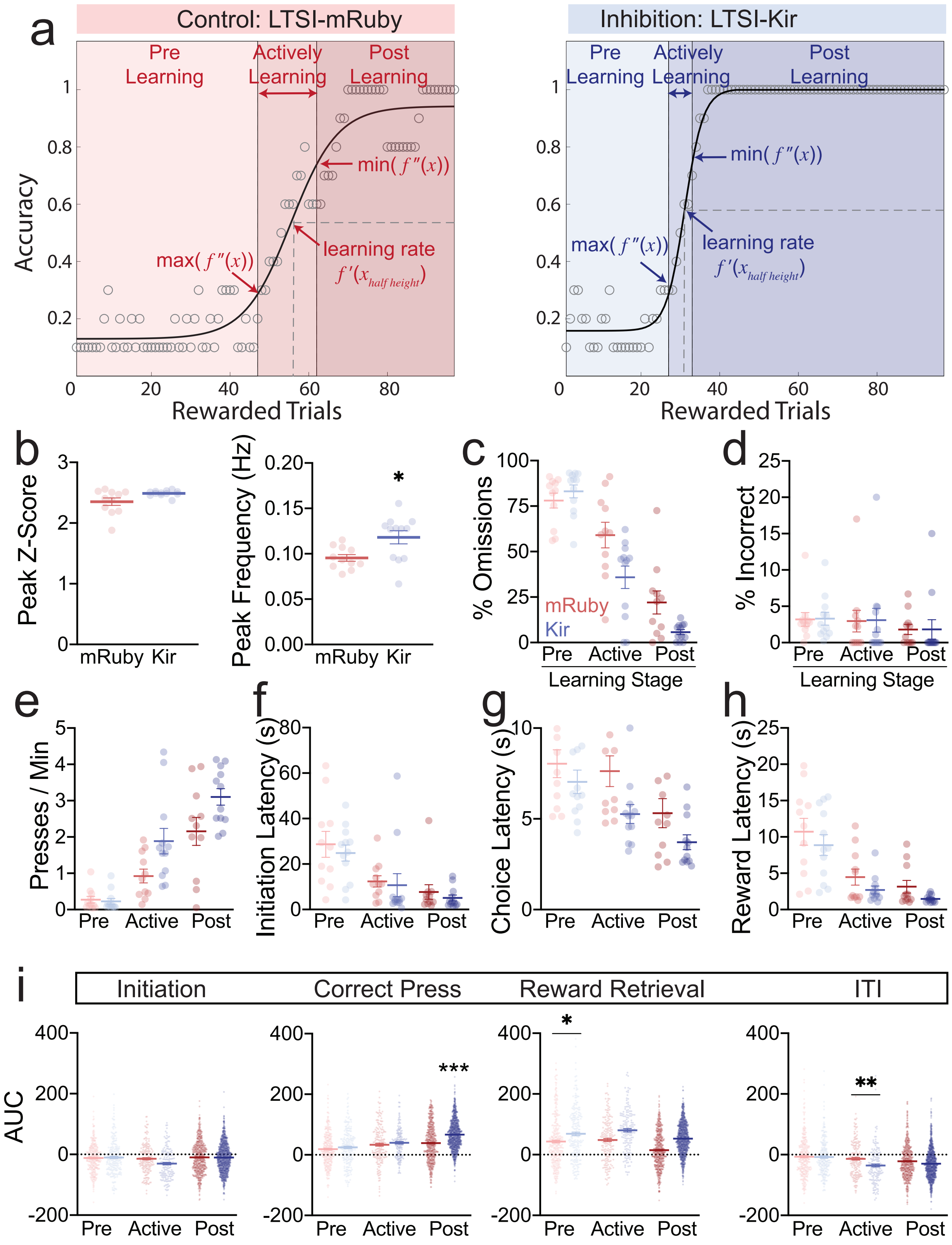


**Supplemental Figure 3. LTSI inhibition amplifies dopamine and accelerates operant learning.** (a) Sample sigmoidal models of learning for a mouse expressing LTSI-mRuby (left) and LTSI-Kir (right). (b) Peak z-score (left) and frequency (right) in the 10min baseline period prior to the operant task of mice expressing LTSI-mRuby (control, n=11) or LTSI-Kir (inhibition, n=12). **p<0.01, ***p<0.001 vs LTSI-mRuby. (c-d) Proportion of (c) omissions (initiations without lever press), and (d) incorrect lever presses in pre-learning, active learning, and post-learning periods. (e) Rate of lever presses across learning stages. (f-h) Latencies to (f) initiate, (g) press a lever, and (h) retrieve reward across learning stages. (i) Total area under the curve (AUC) of the PETH for the 1s before and after behavioral events. **p<0.01, ***p<0.001 vs mRuby control at the same learning stage. Lines in dot plots represent mean ± SEM. See Supplemental Table 1 for detailed statistics.

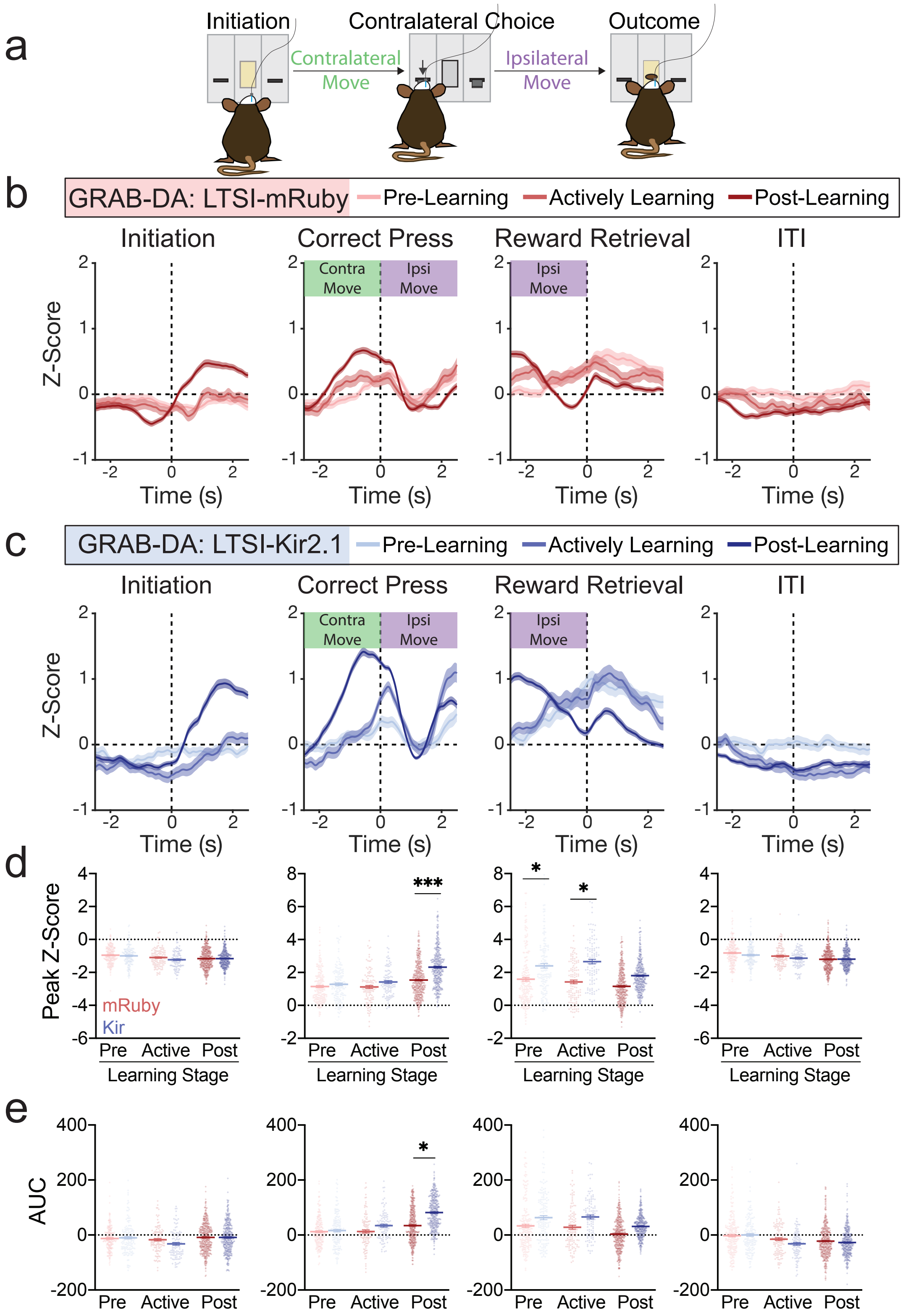


**Supplemental Figure 4. LTSI inhibition amplifies dopamine signals as mice approach a contralateral lever.** (a) As mice press a lever contralateral to the striatal implant, they make a contralateral movement towards the lever followed by an ipsilateral movement towards the reward magazine. (b) Peri-event temporal histograms (PETHs) for initiations, correct presses, reward retrievals, and ITIs of pre-learning, actively learning, and post-learning trials for LTSI-mRuby control mice training to press the lever contralateral to the striatal implant (n=6). (c) PETHs for the same behavioral events for LTSI-Kir2.1 inhibited mice training to press the lever contralateral to the implant (n=7). (d-f) Peak (minima for initiation and ITI, maxima for correct press and reward retrieval) (d) Z-scores, (e) total areas under the curve (AUC) in the window of 1s before and after the behavioral event. *p<0.05, **p<0.01, ***p<0.001 vs mRuby control for same learning stage. Lines in dot plots represent mean ± SEM; all PETH data represented as mean ± SEM. See Supplemental Table 1 for detailed statistics.

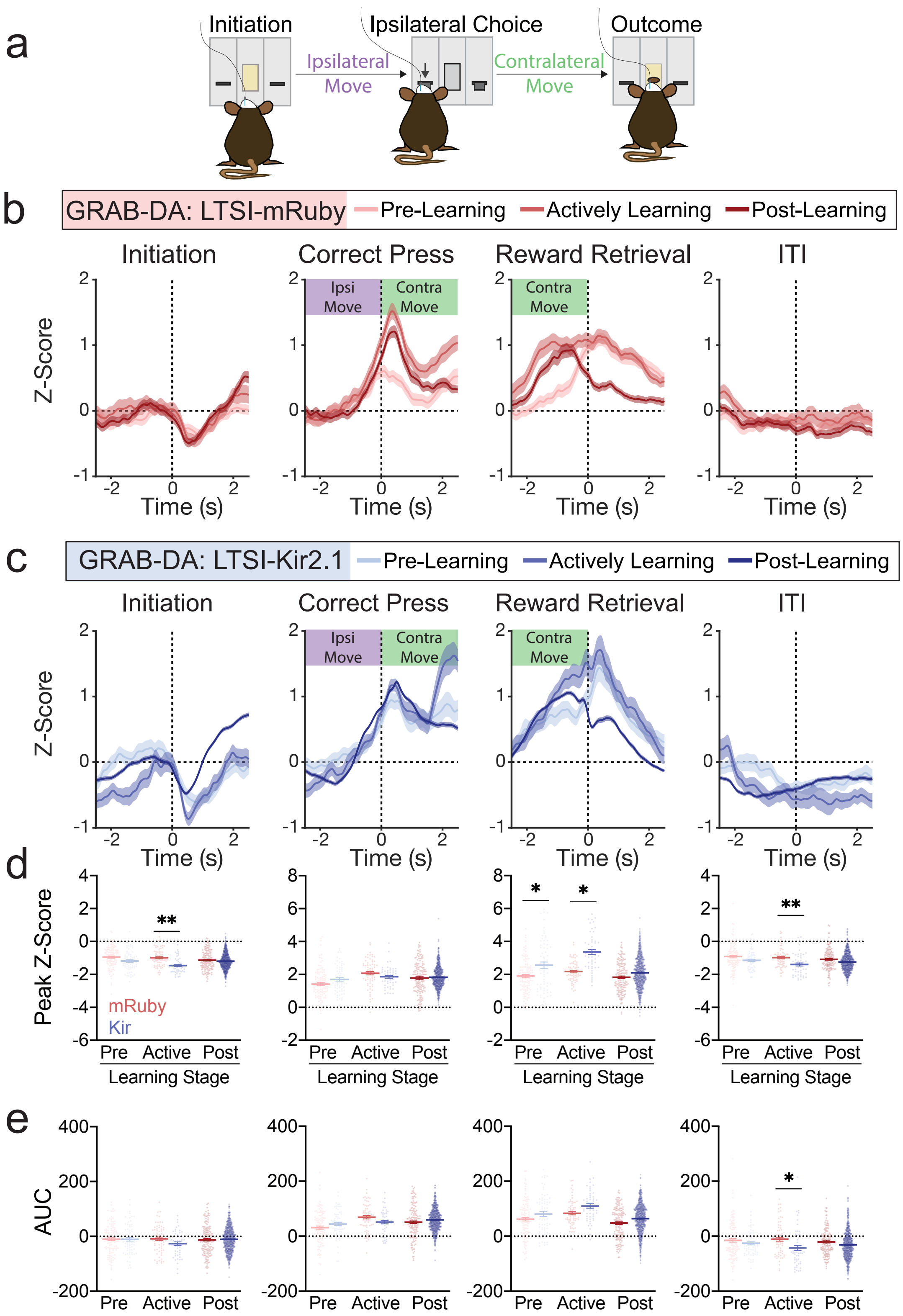


**Supplemental Figure 5. LTSI inhibition amplifies dopamine signals as mice approach a reward contralateral to the striatal implant.** (a) As mice press a lever ipsilateral to the striatal implant, they make an ipsilateral movement towards the lever followed by a contralateral movement towards the reward magazine. (b) Peri-event temporal histograms (PETHs) for initiations, correct presses, reward retrievals, and ITIs of pre-learning, actively learning, and post-learning trials for LTSI-mRuby control mice training to press the lever ipsilateral to the striatal implant (n=5). (c) PETHs for the same behavioral events for LTSI-Kir2.1 inhibited mice training to press the lever ipsilateral to the implant (n=5). (d-f) Peak (minima for initiation and ITI, maxima for correct press and reward retrieval) (d) Z-scores, and (e) total areas under the curve (AUC) in the window of 1s before and after the behavioral event. *p<0.05, **p<0.01, ***p<0.001 vs mRuby for same learning stage. Lines in dot plots represent means ± SEM; all PETH data represented as mean ± SEM. See Supplemental Table 1 for detailed statistics.

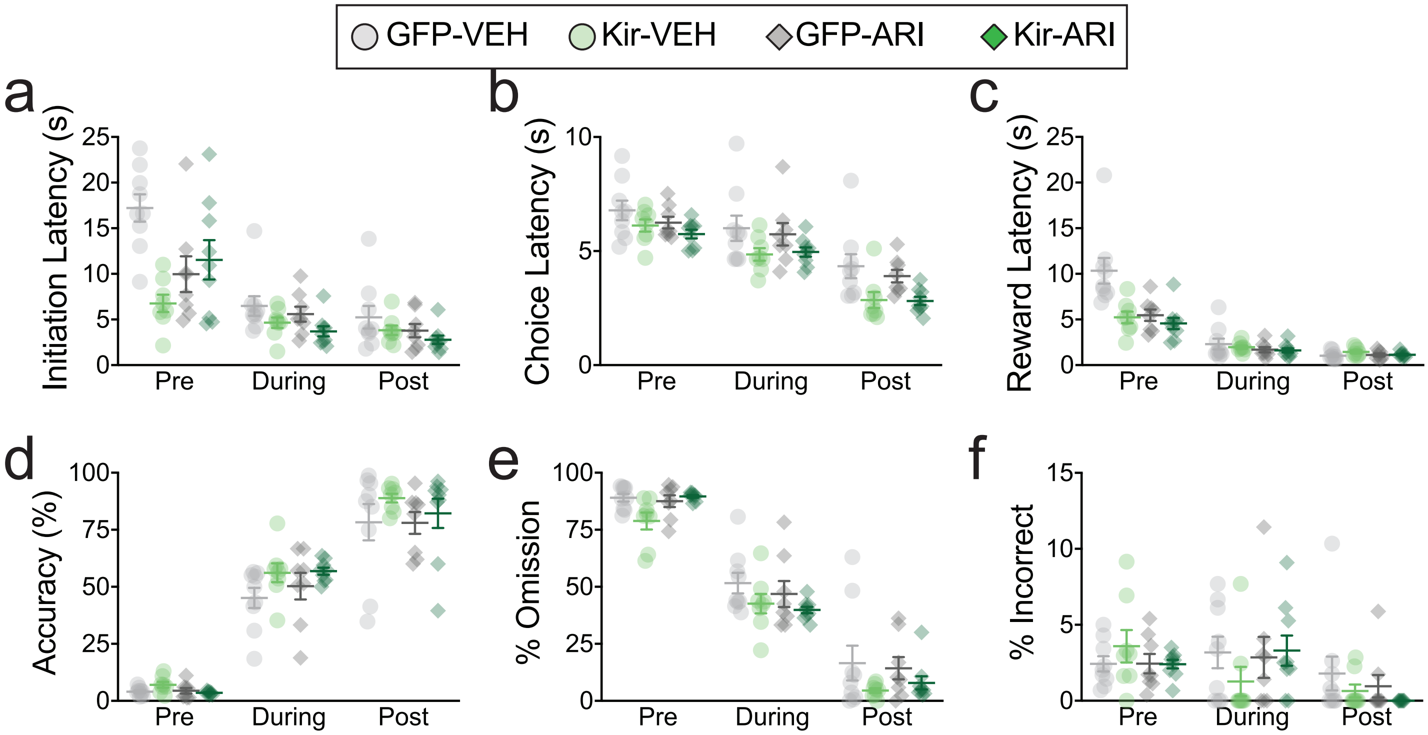


**Supplemental Figure 6. Intra-striatal dopamine D2 partial agonism prevents effects of LTSI inhibition on learning.** (a-c) Latencies to (a) initiate, (b) lever press, and (c) retrieve reward across learning stages. (d-f) Proportions of (d) correct presses, (e) omitted responses, and (f) incorrect lever presses across learning stages. All individual data shown, with bars representing mean ± standard error. See Supplemental Table 1 for detailed statistics.
